# An essential role for glial cells in postingestive nutrient sensing

**DOI:** 10.1101/2024.07.26.605306

**Authors:** Divita Kulshrestha, Charlotte Koch, Majid Khalilov, Stefanie Schirmeier

## Abstract

Behavioral flexibility is an essential trait for survival of any organism. For example, choosing an appropriate food source is vital. To do so, the food’s nutritive value needs to be evaluated. Several studies have demonstrated that *Drosophila melanogaster* larvae and adults, as mammals, are able to distinguish between nutritious and non-nutritious carbohydrates independent of their taste. Several groups of neurons have been implicated in postprandial sugar sensing in adult flies (Dus et al., 2011, 2015; Miyamoto et al., 2012; Park et al., 2016; Musso et al., 2023). In larvae, neurons expressing Gr43a, a fructose receptor, have been shown to be implicated in postingestive glucose sensing (Mishra et al., 2013). How can a fructose sensor mediate glucose sensing? We show that postingestive glucose, and also sorbitol, sensing involves carbohydrate conversion into fructose via the polyol pathway in glial cells. Glia-derived fructose is subsequently sensed via Gr43a expressed in neurons, which leads to behavioral adaptation. Thus, in postingestive nutrient sensing, the glial cells play a central role in information processing and regulation of behavior.

## Introduction

Choosing appropriate food sources is vital to the success of any organism. Food composition is evaluated in two ways, preingestively, via taste and smell, and postingestively. Feeding is mainly regulated via gustatory cues, since they allow assessment of palatability and nutrient content. However, it is modulated by postingestive mechanism that allow sensing of the nutritive value of ingested food and thereby maintaining homeostasis (Sung et al., 2023). Amino acids, lipids and carbohydrates are the macronutrients essential for an organism. Carbohydrates are a major source of energy. Nutritive carbohydrates provide a feeding stimulus and are sensed via their sweet taste, but also get postingestively evaluated for their caloric value (Margolskee, 2002). Postingestive carbohydrate sensing takes place in different organs. In mammals, the pancreas, the gut-brain axis and the brain are central to glucose sensing and feeding regulation (Rosario et al., 2016; Tan et al., 2020; Yoon and Diano, 2021).

Postingestive nutrient sensing is not specific to mammals, but also exists in other organisms, like insects. Here, postingestive evaluation of the nutritive value of carbohydrates takes place in different organs including the gut, the fatbody and the brain. The brain, thus, has a dual role in regulating food choice and feeding behavior: It acts as a nutrient sensor and is responsible for the behavioral output. In the adult fly, internal nutrient sensors have been shown to play a crucial role in discriminating between nutritive and non-nutritive carbohydrates. Several groups of central neurons have been shown to sense carbohydrates and regulate feeding behavior: the diuretic hormone 44 (Dh44)- expressing neurons, SLC5A11-expressing R4 neurons and Gr43a-positive neurons (Dus et al., 2011, 2015; Miyamoto et al., 2012; Park et al., 2016; Musso et al., 2023). Dh44-expressing neurons and SLC5A11-expressing neurons are thought to sense glucose, while Gr43a is a fructose sensor (Dus et al., 2011, 2015; Park et al., 2016; Musso et al., 2023). Larval Drosophila can sense carbohydrates as well. In contrast to adults, the larva only expresses one gustatory carbohydrate receptor, Gr43a, which is fine-tuned to fructose (Mishra et al., 2013; Fujii et al., 2015). Gr43a-expression in the pharyngeal taste neurons allows the larva to taste fructose. Interestingly, the larva is also able to sense other nutritive carbohydrates, like glucose or sorbitol, even though it cannot taste them (Mishra et al., 2013). Gr43a-expressing neurons in the central nervous system (CNS) have been shown to be essential for this postingestive sensing of carbohydrates (Mishra et al., 2013). However, it remains unclear how the fructose receptor, Gr43a, expressed in the CNS can mediate glucose sensing. SLC5A11 has also been implicated in glucose sensing in Drosophila larvae, but seems to be regulated by insulin and mainly needed to sense hypoglycemia (Ugrankar et al., 2018).

Here, we show that postingestive glucose sensing in the larva indeed takes place inside the CNS, since efficient carbohydrate uptake into the nervous system is a prerequisite. Inside the nervous system, glial cells sense glucose concentrations via aldose reductases and metabolize excess glucose to fructose, which in turn is sensed by Gr43a expressed in neurons. The Gr43a-expressing neurons use Corazonin as a neurotransmitter to regulate feeding behavior. Thus, we highlight the role of glial cells as carbohydrate sensor in the larval CNS.

## Results

### A frustrated total internal reflection-based imaging setup can be used to study postingestive nutrient sensing-dependent food choice behavior

We employed a frustrated total internal reflection (FTIR)-based imaging setup (FIM; (Risse et al., 2013, 2017) to study the mechanism of Gr43a-mediated glucose and sorbitol sensing in larvae. This setup has previously been shown to reliably track larval fructose-dependent food choice behavior (Bittern et al., 2022). To test our setup, we aimed to reproduce the data on glucose and sorbitol dependent food choice published by Mishra and colleagues previously (Mishra et al., 2013). We were able to reproduced the immediate or delayed preference of larvae presented with either fructose, sorbitol or glucose (Fig. 1). As expected, larvae given a choice between plain agar and fructose choose fructose fast (Fig. 1 A,B). Sorbitol and glucose preference are delayed compared to fructose preference (Fig. 1C-F). In accordance with the published data (Mishra et al., 2013), preference for all carbohydrates depends on Gr43a-expression, since panneuronal RNA interference-mediated knockdown of Gr43a abolishes the ability of larvae to choose carbohydrate agar over plain agar (Fig. 1). Thus, our FIM-based imaging method can be applied to study larval food choice behavior. Since sorbitol and glucose preference develop with a delay (depends on the experiment, in average about 8 min for sorbitol and about 12 minutes for glucose) a significant difference between Gr43a-KD animals and controls is just seen after these orientation phases (Fig. 1D,F).

**Figure 1:**
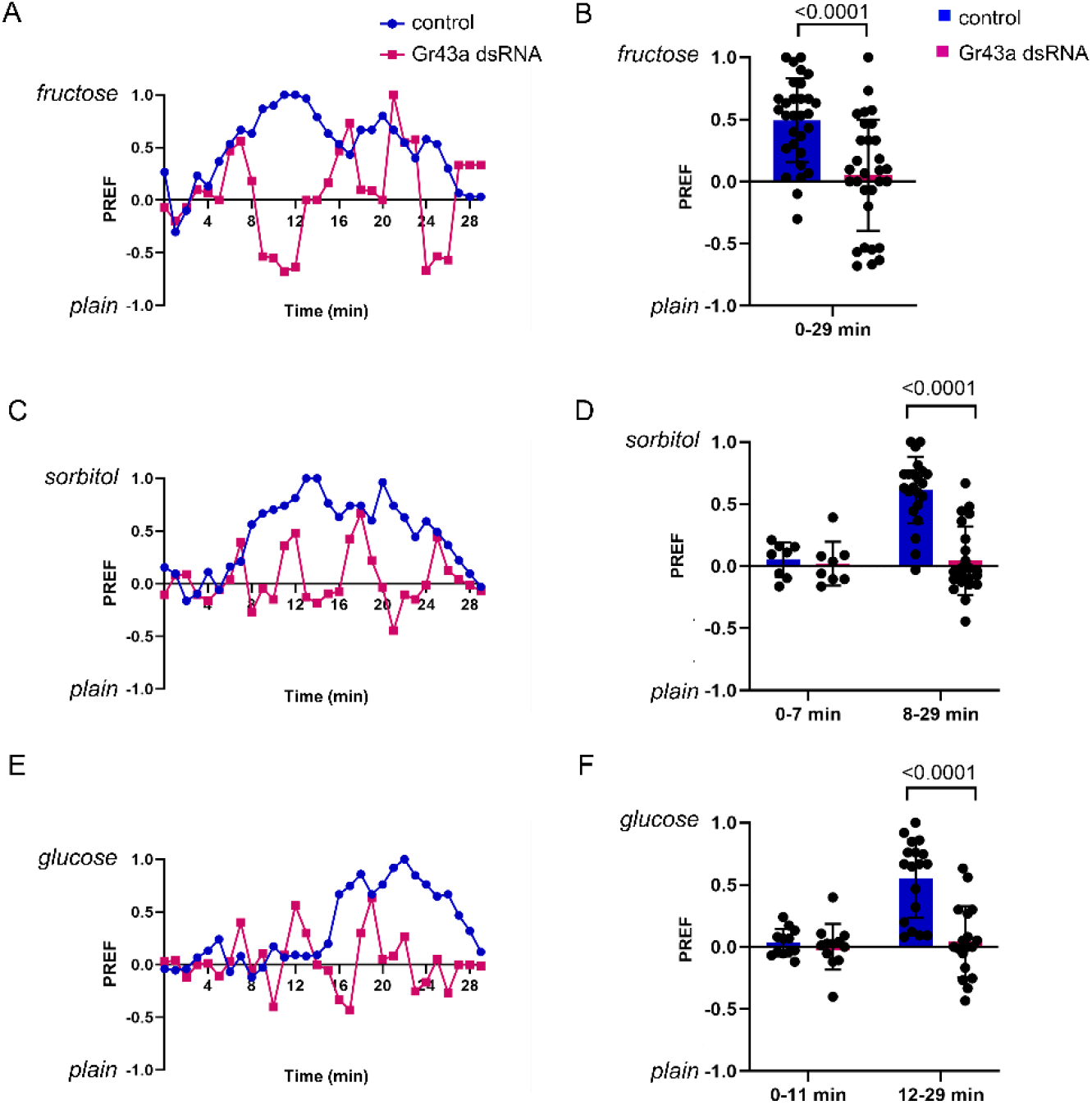
A FIM-based tracking setup can be used to study postingestive nutrient sensing in Drosophila larvae. As published previously, Gr43a expression is essential for carbohydrate sensing in Drosophila larvae. A,B) Panneuronal knockdown of Gr43a abolishes fructose preference. A) Example preference indices over time. B) Quantification of fructose preference. Preference development is fast. N=4, n=80 C,D) Panneuronal knockdown of Gr43a abolishes postingestive sorbitol preference. C) Example preference indices over time. D) Quantification of sorbitol preference. During the orientation phase (min 1-7) no difference between the two genotypes can be seen, while in the preference phase (min 8-29) the control animals clearly prefer sorbitol-containing agar over plain agar. N=4, n=80 E,F) Panneuronal knockdown of Gr43a abolishes postingestive glucose preference. E) Example preference indices over time. F) Quantification of glucose preference. During the orientation phase (min 1-11) no difference between the two genotypes can be seen, while in the preference phase (min 12-29) the control animals clearly prefer glucose-containing agar over plain agar. N=4, n=80. Comparison of two groups was performed using Mann-Whitney U test.

### Dh44 positive neurons are not implicates in postingestive glucose or sorbitol sensing in the larva

In adult Drosophila several CNS neuronal populations have been implicated in postingestive glucose sensing (Dus et al., 2011, 2013, 2015; Miyamoto et al., 2012). Besides Gr43a-expressing neurons, also SLC5A11-expressing neurons and Dh44 (diuretic hormone 44)-expressing neurons have been shown to be essential for glucose sensing. Dh44 is also expressed in the larval CNS (Fig. 2A, Zandawala et al., 2018), but its role in carbohydrate sensing has not been analyzed. Thus, we knocked down Dh44 in all neurons or ablated Dh44 neurons (using Dh44-Gal4 driven UAS-hid; hid is an activator of apoptosis) and analyzed larval ability to sense glucose (Fig. 2). In contrast to Gr43a-knockdown, panneuronal Dh44-knockdown does not have any effect on postingestive glucose sensing (Fig. 2B) and even ablation of Dh44-positive neurons does not show any effect (Fig.2C). In contrast, ablation of Gr43a-neurons abolishes glucose preference efficiently. Thus, Dh44-expressing neurons do not seem to be implicated in postingestive glucose sensing in Drosophila larvae.

**Figure 2:**
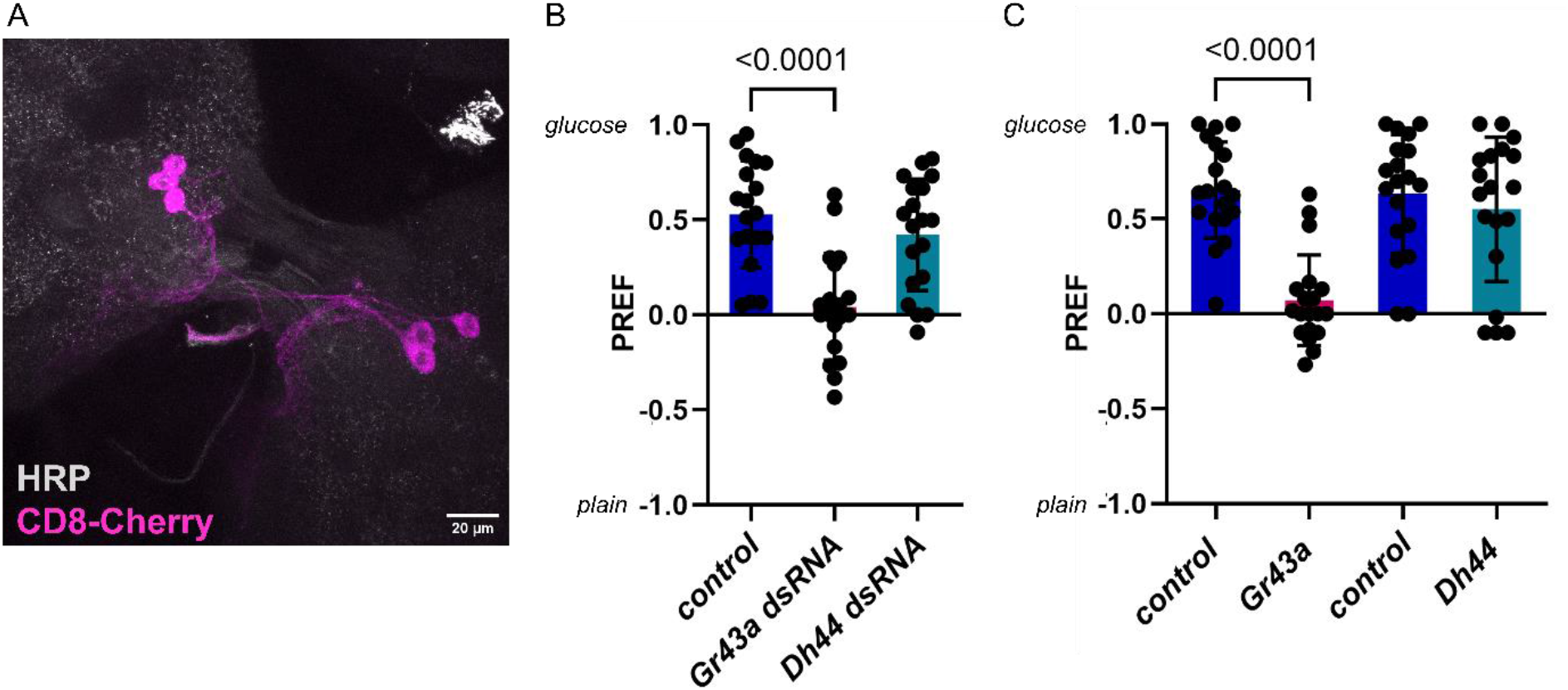
Gr43a-expressing neurons, but not Dh44-expressing neurons are implicated in postingestive glucose sensing. A) Dh44 is expressed in 6 cells in the larval central brain. CD8-Cherry expression is driven using Dh44-Gal4 (magenta). Neuronal membranes are stained using α-HRP (grey). B) Panneuronal knockdown of Gr43a abolishes postingestive glucose preference, while Dh44 knockdown has no influence on glucose preference. Mean preference indices of min 12-29 are shown. N=4, n=80 C) Ablation of Gr43a-expressing neurons using Gr43a-lexA driven lexAop-hid abolishes glucose preference, while ablation of Dh44-expressing neurons using Dh44-Gal4 driven UAS-hid does not affect food choice. Control animals express UAS-mCD8GFP under the control of the respective driver. Mean preference indices of min 12-29 are shown. N=4, n=80. Comparison of two groups was performed using Mann-Whitney U test.

### Efficient carbohydrate transport over the BBB is essential for development of glucose preference

As Gr43a-positive neurons in the CNS are essential for fructose sensing (Mishra et al., 2013), the question arises if carbohydrates need to be taken up into the nervous system to be sensed. To answer this question, we made use of pippin knockdown and knockout animals, that we had studied earlier (McMullen et al., 2021). When pippin is knocked down in either the perineurial or subperineurial glial cells of the blood-brain barrier (BBB), glucose uptake into the respective cell type is severely reduced (McMullen et al., 2021). Pippin mutant animals are partially compensated and display wildtypic carbohydrate uptake into the perineurial glial cells, but reduced carbohydrate uptake into subperineurial glial cells. Thus, panglial knockdown of pippin induces pupal lethality, while mutants are viable (McMullen et al., 2021). We tested both pippin mutants and panglial pippin knockdown animals for their capacity to choose different carbohydrates (Figure 3). Pippin mutant larvae are able to choose glucose over plain agar, but show a delay in preference development (Figure 3A). Panglial knockdown of pippin, which shows stronger impairment of carbohydrate transport into the nervous system, abolishes glucose preference completely (Figure 3C). Interestingly, sorbitol preference is not affected by loss of pippin, indicating that sorbitol is transported into the brain independently of pippin (Figure 3B,D). Nonetheless, this data suggests that efficient carbohydrate transport into the nervous system is essential for postingestive carbohydrate sensing.

**Figure 3:**
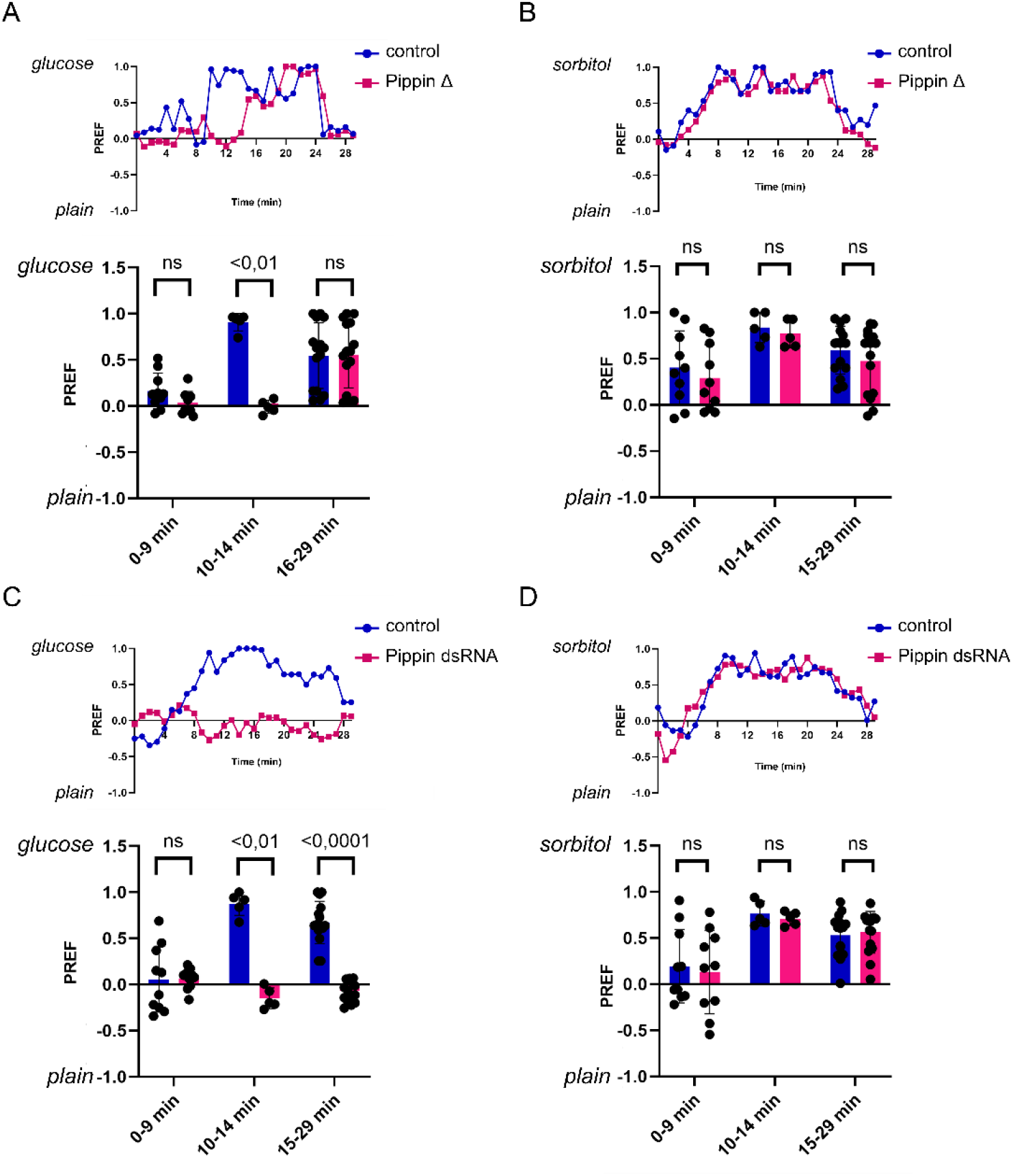
Efficient glucose transport over the BBB is essential for postingestive glucose sensing. A) Pippin mutant animals show develop glucose preference later than control animals. Preference indices are shown over time (upper panel) and as means for different time intervals. In the orientation phase both genotypes do not display any preference. Control animals already display a preference for glucose in the 10-14 min interval, while pippin mutants just display a preference from 15 min onwards. N=4, n=80. B) Pippin mutants display wildtypic postingestive sorbitol preference. Preference indices are shown over time (upper panel) and as means for different time intervals (lower panel). N=4, n=80. C) Panglial *pippin* knockdown abolishes glucose preference. Preference indices are shown over time (upper panel) and as means for different time intervals. N=4, n=80. D) Panglial *pippin* knockdown animals display wildtypic postingestive sorbitol preference. Preference indices are shown over time (upper panel) and as means for different time intervals (lower panel). N=4, n=80. Comparison of two groups was performed using Mann-Whitney U test.

### Conversion of glucose/sorbitol to fructose via the polyol pathway in glia is essential for postingestive sensing

Since Gr43a is fine-tuned fructose sensor (Miyamoto et al., 2012), we wondered how Gr43a can be implicated in glucose and sorbitol sensing. We hypothesized that glucose and sorbitol could be converted into fructose via the polyol pathway. This pathway converts Glucose into Fructose via Sorbitol and has been shown to act as a fructose sensing mechanism in other tissues (Figure 4, Sano et al., 2022). In the polyol pathway, glucose is first converted into sorbitol by an aldose reductase (AR) and then into fructose via sorbitol dehydrogenase (Sodh). The Drosophila genome encodes eight potential ARs (CG6084, CG2747, CG10863, CG6083, CG9436, CG12766, CG10638, CG40064), two of which show high expression in larvae (FlyAtlas Anatomical Expression Data and modENCODE), CG6084 (Aldo-keto reductase 1B, Akr1B) and CG10863 (Aldo-keto reductase 2E3, Akr2E3). To analyze the role of these two ARs, we knocked both down using either a panglial (repo) or a panneuronal (elav) driver (figure 4). Animals with a glial knockdown of either AR display a strong impairment of glucose sensing, while sorbitol sensing is not affected, as could be expected (Fig. 4 B, D). Double knockdown of both ARs in glial cells (AR dsRNA) abolished glucose preference completely. Neuronal knockdowns, however, did not induce any abnormal phenotypes (Fig4 C, E). Thus, glucose seems to be converted to sorbitol in glial cells via Akr1B and Akr2E3. The Drosophila genome encodes two sorbitol dehydrogenases, Sodh1 and Sodh2. Only Sodh2 (CG4649) is broadly expressed in larval tissues. Glial knockdown of Sodh2 strongly impairs development of both glucose and sorbitol preference (Fig. 4B, D). Neuronal knockdown again does not induce any abnormal phenotypes (Fig. 4C, E). Thus, a functional polyol pathway in glial cells is essential for postingestive glucose and sorbitol sensing.

**Figure 4:**
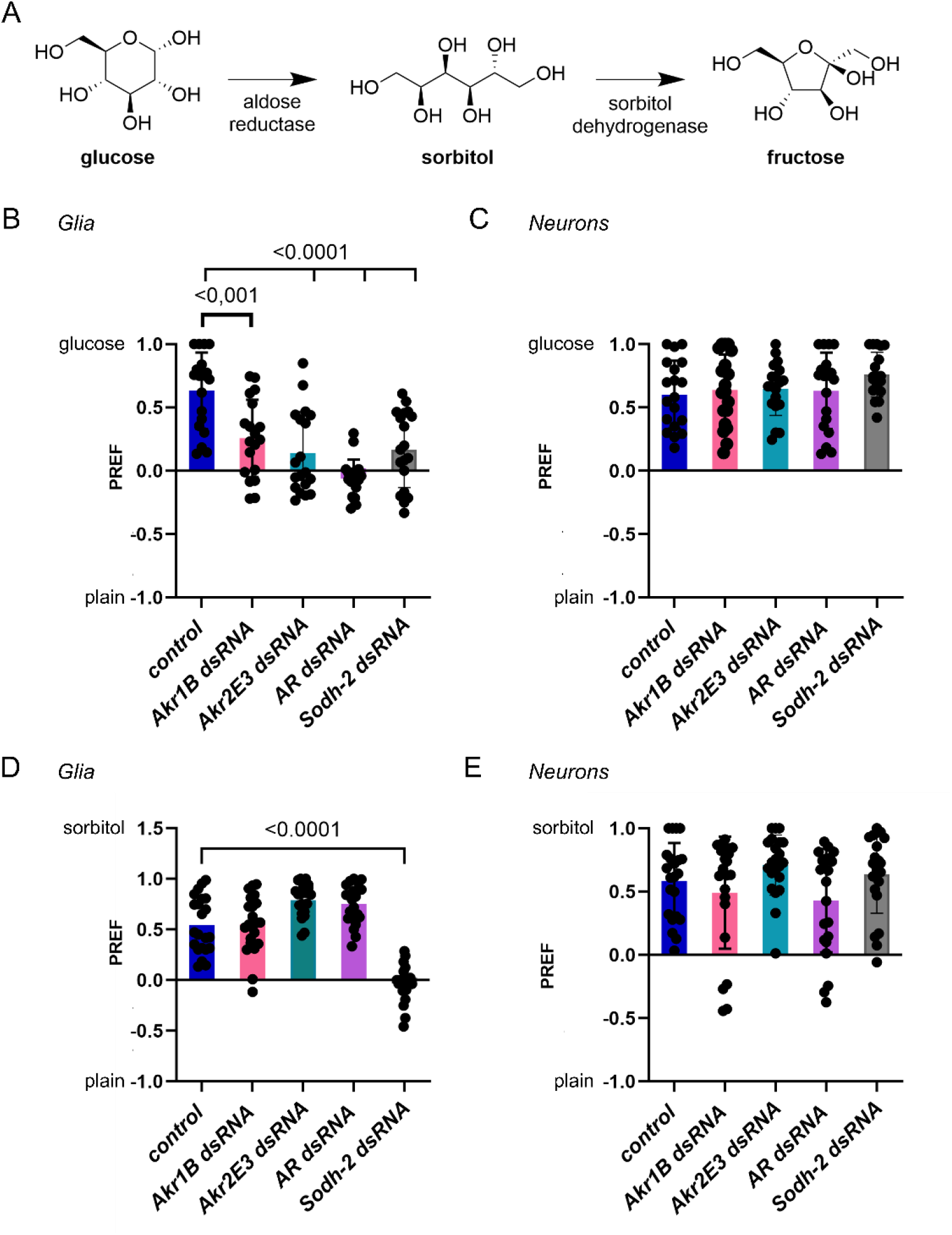
Glial carbohydrate processing via the polyol pathway is essential for postingestive glucose and sorbitol sensing. A) Schematic representation of the polyol pathway. Glucose gets reduced by aldose reductases (AR) to sorbitol. Sorbitol is the substrate for sorbitol dehydrogenase (Sodh) that produces fructose. B) Panglial knockdown of ARs Akr1B or Akr2E3 significantly reduces glucose preference. Double knockdown of both ARs (AR dsRNA) abolishes glucose preference. Panglial knockdown of Sodh2 also reduces glucose preference significantly. Mean preference indices of min 12-29 are shown. N=4, n=80. C) Panneuronal knockdown of either ARs or Sodh2 does not influence glucose preference. Mean preference indices of min 12-29 are shown. N=4, n=80. D) Panglial knockdown of ARs, Akr1B, Akr2E3 or both, does not influence sorbitol preference. Panglial knockdown of Sodh2 reduces sorbitol preference significantly. Mean preference indices of min 8-29 are shown. N=4, n=80. C) Panneuronal knockdown of either ARs or Sodh2 does not influence sorbitol preference. Mean preference indices of min 8-29 are shown. N=4, n=80. Comparison of two groups was performed using Mann-Whitney U test.

### Cortex as well as astrocyte-like glia are implicated in carbohydrate conversion

To understand which glial subtypes convert glucose and sorbitol to fructose, we performed glial subtype-specific knockdown experiments. We either knocked down both ARs or Sodh2 using the following drivers: 9137-Gal4 (subperineurial and perineurial glial cells = surface glial cells (SG); Desalvo et al., 2014), R83E12-Gal4 (ensheathing glial cells; Li et al., 2014), alrm-Gal4 (astrocyte-like glial cells; Muthukumar et al., 2014), and R55B12 (Cortex glial cells; Li et al., 2014). Knockdown of either both ARs or Sodh2 in the surface glia, ensheathing glia, astrocyte-like glia or cortex glia alone induced no significant changes in glucose or sorbitol preference (Figure 5). However, cortex glial knockdown of ARs displayed a tendency towards lower preference. When we combined drivers for cortex glial and astrocyte-like glial expression, AR knockdown and Sodh2 knockdown abolished glucose or sorbitol preference, respectively (Figure 5), indicating that both cell types participate in producing fructose from glucose or sorbitol.

**Figure 5:**
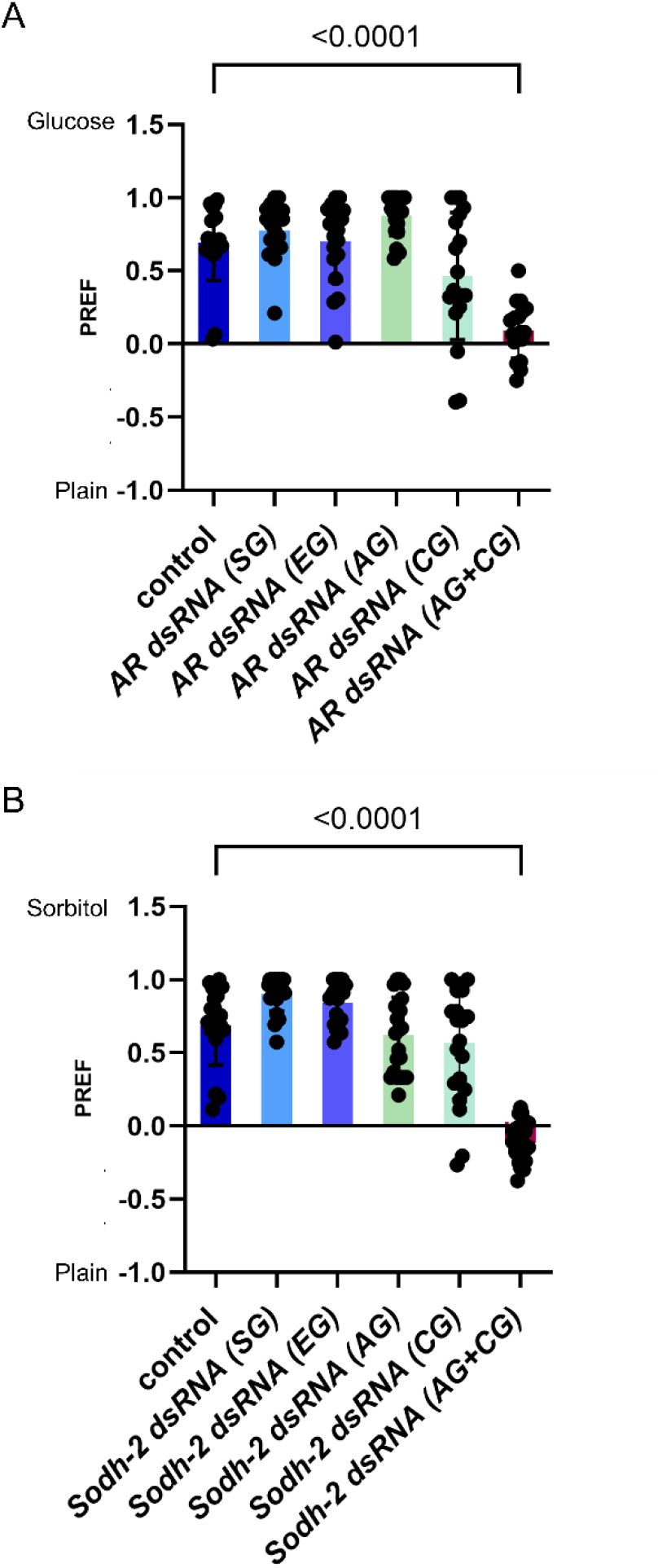
Astrocyte-like and cortex glia produce fructose from glucose. A) Knockdown of ARs in different glial subtypes. Only knockdown of ARs in both the cortex glia and the astrocyte-like glia simultaneously abolishes glucose preference. Mean preference indices for the time interval 12-29 min are shown. N=4, n=80. B) Knockdown of Sodh2 in different glial subtypes. Only knockdown of Sodh2 in both the cortex glia and the astrocyte-like glia simultaneously abolishes sorbitol preference. Mean preference indices for the time interval 8-29 min are shown. N=4, n=80. SG: surface glia (perineurial and subperineurial glia, 9137-Gal4); EG: ensheathing glia (nrv2-Gal4); AG: astrocyte-like glia (alrm-Gal4); CG: cortex glia (GMR55B12-Gal4). Comparison of two groups was performed using Mann-Whitney U test.

### Larval Gr43a neurons signal to downstream neurons via the neuropeptide Corazonin

Since Gr43a neurons are regulating food choice behavior upon fructose signaling from the glial cells, it is of interest to understand how they signal to downstream target neurons. It has been previously reported that some Gr43a-positive neurons express the neuropeptide Corazonin (Crz) (Fig. 6A, Miyamoto and Amrein, 2014). Crz is an orthologue of mammalian gonadotropin releasing hormone. Amongst others, it has been implicated in regulating Drosophila alcohol resistance, stress response, food intake and metabolism (Kapan et al., 2012; McClure and Heberlein, 2013; Kubrak et al., 2016, 2022). To understand the role of Crz in Gr43a-dependent food choice behavior, we knocked down Crz in Gr43a-expressing neurons specifically. Crz-KD severely reduces glucose as well as sorbitol preference (Fig. 6B). To further validate the role of Crz-signaling we knocked down the Crz receptor (CrzR) in all neurons. Loss of CrzR phenocopies Crz KD in Gr43a-positive neurons (Fig. 6C), indicating that indeed Gr43a-neurons are signaling to downstream neurons via Crz. As expected this signaling is not needed to develop an immediate fructose preference, which can be established independent of the central Gr43a-expressing neurons (Fig 6C; Mishra et al., 2013)).

**Figure 6:**
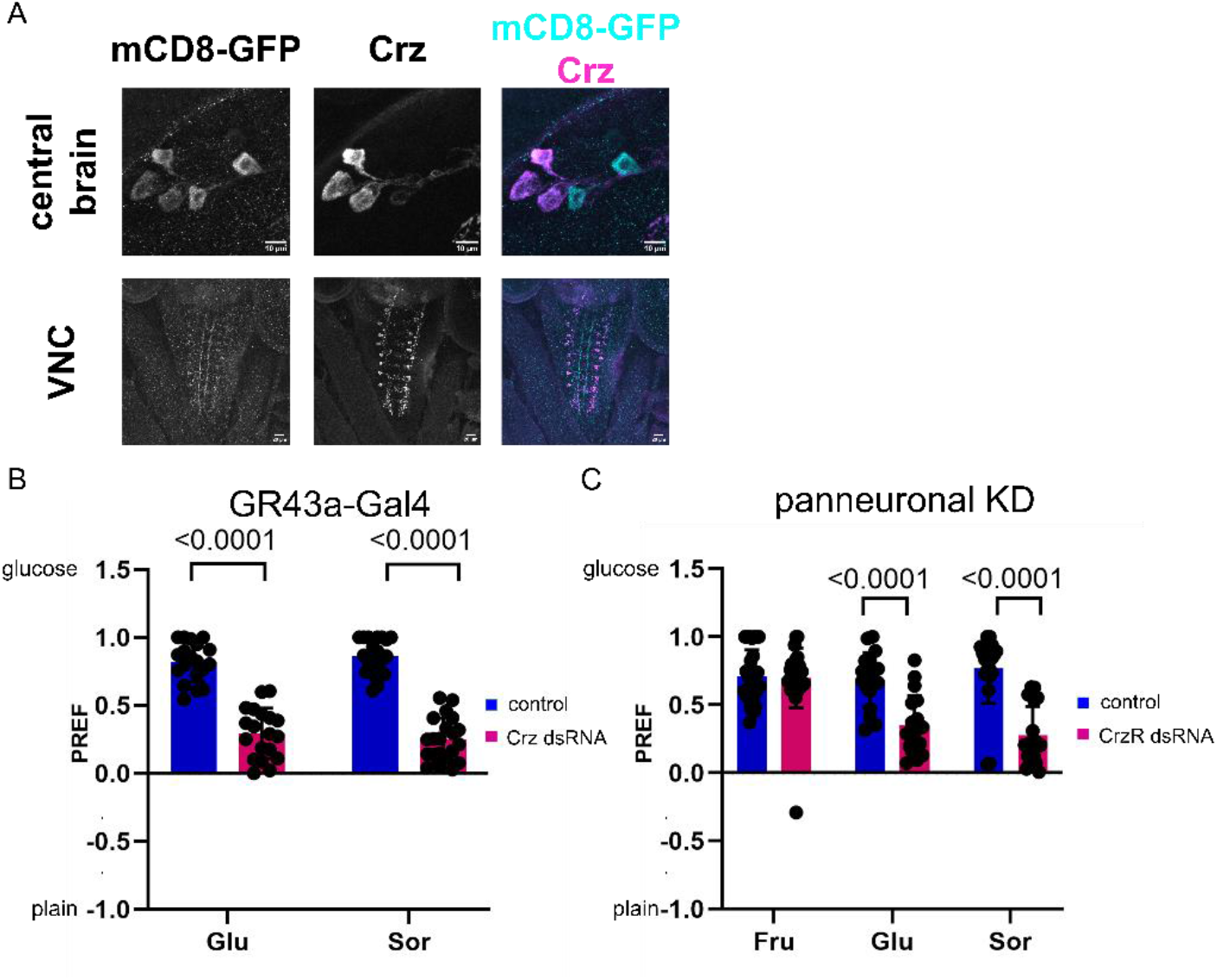
GR43a expressing-neurons signal via Crz. A) Immunofluorescence staining of L3 larval brains expressing mCD8-GFP (cyan) under GR43a control (Gr43a-lexA) for Crz (magenta). Upper panels show GR43a positive cells (mCD8-GFP) in the central brain, lower panels show GR43a-positive neurons (mCD8-GFP) in the ventral nerve cord. B) Gr43a-expressing neuron-specific knockdown of Crz significantly reduces preference for glucose and sorbitol. Mean preference indices for the time interval 12-29 min (glucose) or 8-29 min (Sorbitol) are shown. N=4, n=80. C) panneuronal knockdown of CrzR significantly reduces preference for glucose and sorbitol. Fructose preference is wild typic. Mean preference indices for the time interval 1-29 min (fructose), 12-29 min (glucose) or 8-29 min (Sorbitol) are shown. N=4, n=80. Comparison of two groups was performed using Mann-Whitney U test.

## Discussion

For a long time, the polyol pathway has been thought to not play a major role in healthy organisms, since the affinity of aldose reductases to glucose is rather low, especially compared to other glucose-processing enzymes such as hexokinase. Nevertheless, it has been suggested to be activated upon hyperglycemia and to be the cause for many complications in diabetic patients since a long time (Gabbay, 1973). Recently, it has also come into focus as a metastasis promoting factor (Wu et al., 2017; Schwab et al., 2018; Kang et al., 2024). In addition, it has been suggested to serve as a conserved cellular glucose sensing mechanism active in the Drosophila fat body and mammalian liver (Sano et al., 2022). Here, we show that it also acts as a cellular glucose sensing mechanism in the Drosophila brain. Since aldose reductases and sorbitol dehydrogenases seem to be expressed broadly (Tang et al., 2012), the polyol pathway could potentially serve as a cellular glucose sensor also in other cell types.

Originally thought to have only structural roles, glial cells are nowadays known to serve divers functions in the nervous system. Their roles range from functions in nervous system development to modulating synaptic communication, plasticity, homeostasis, and activity of neuronal networks (Allen and Lyons, 2018). However, it has been common sense that the sensory cells of the nervous system are neurons. Here, we show that the glial cells in the central nervous system of Drosophila larvae sense carbohydrate concentration changes using the polyol pathway as a molecular sensor. Via this pathway high concentrations of carbohydrates are sensed and used to produce fructose. This glia-derived fructose then activates GR43a-expressing neurons to convey the signal and induce behavioral changes. A sensory role has also been suggested for specific peripheral glial cells in C. elegans. Here, AMsh glia detect repulsive odorants and induce olfactory adaptation (Duan et al., 2020). Taken together, we likely will have to expand the vast catalogue of glial functions by the term “sensory functions”.

## Methods

### Fly stocks and husbandry

Fly stocks were kept at room temperature or 25°C on standard cornmeal medium. The stocks were used: Dh44-Gal4 (BDSC 51987), Gr43a-Gal4 (BDSC 57636), Gr43a-lexA (BDSC 93446), repo-Gal4;repo-Gal4 (Sepp and Auld, 1999; Lee and Jones, 2005), elav-Gal4;;elav-Gal4 (BDSC 458, BDSC 8760), nrv2-Gal4 (BDSC 6797), GMR55B12-Gal4 (BDSC 39103), 9-137-Gal4 (Desalvo et al., 2014), alrm-G4 (Doherty et al., 2009), UAS-hid (S. Luschnig), lexAop-hid (C. Klämbt), UAS-mCD8GFP (BDSC), UAS-Dh44-dsRNA(BDAC 25804), UAS-Gr43a-dsRNA (BDSC 64881), UAS-AkrB1-dsRNA (BDSC 62219), UAS-Akr2E3-dsRNA (BDSC 34878 or VDRC 48619), UAS-Sodh2-dsRNA (BDSC 53353), UAS-mCherry-dsRNA (BDSC 35785) was used as control dsRNA.

### Two-choice-assay

Food choice behavior of fed third instar feeding larvae was recorded as previously described (Bittern et al., 2022). Ten larvae were used per arena. Recording was done at room temperature (24-26°C) in the absence of light. Data analysis has been performed using FIMTrack and a custom-written MATLAB script as described previously (Bittern et al., 2022). The preference index is calculated as follows: sugar^Preference index^ = (#larvae^sugar^ – #larvae^plain agar^) / total #larvae; it gives the value 0 when no preference is present, +1 when all animals are on the sugar side, or -1 when all animals are on the plain agar side.

### Immunohistochemistry

Feeding third instar larvae were washed and collected in ice-cold water. Open book preparations or larval filets were prepared using needles and forceps in ice-cold 1x PBS. Larval preparations were stained following standard protocols. Samples were imaged using the Zeiss LSM800 confocal microscope, using a plan apochromat DIC III 40/1.3 oil objective at 1024×1024 resolution at 0.65μm z-step size. All images were processed with ZenBlue 2012 (Carl Zeiss AG) and Fiji. The following antibodies were used: anti-Crz (rabbit, 1:1000, Jan Veenstra), anti-GFP (chicken, 1:500, Abcam), anti-mCherry (rat, 1:1000, Invitrogen), anti-HRP DyLight649 (1:250, Dianova), secondary antibodies form Invitrogen and Life Technologies Europe were used.

## Acknowledgement

The work was supported by a grant of the DFG to SS (SCHI1380/5-1).

## Notes

### Competing Interest Statement

The authors have declared no competing interest.

